# The Nucleosome Remodelling and Deacetylation complex restricts Mediator access to enhancers to control transcription

**DOI:** 10.1101/103192

**Authors:** Maria Xenophontos, Nicola Reynolds, Sarah Gharbi, Ewan Johnstone, Jason Signolet, Robin Floyd, Meryem Ralser, Susanne Bornelöv, Sabine Dietmann, Remco Loos, Paul Bertone, Brian Hendrich

## Abstract

A number of different chromatin remodelling complexes in mammalian cells are implicated in the control of gene expression. The genetic requirements for many such complex components have been described, and the biochemical activities of complex components characterised in vitro, yet the molecular mechanisms by which these biochemical activities impact transcriptional regulation in vivo remain ill-defined. Using an inducible system with fine temporal resolution, we show that the Nucleosome Remodelling and Deacetylation (NuRD) complex directly regulates chromatin architecture at enhancer regions in ES cells, in turn influencing the activity of RNA polymerase II via Mediator. Through this mechanism NuRD restricts Mediator access to enhancer chromatin during lineage commitment, thereby enabling appropriate transcriptional regulation. In contrast, acetylation levels of histone H3 lysine 27 are not immediately impacted by NuRD activity, correlating with transcriptional response only after expression levels have changed. These findings provide a detailed, molecular picture of genome-wide modulation of lineage-specific transcription by an abundant chromatin remodelling complex.

## Introduction

Cellular identity is fundamentally determined by the cohort of genes activated and repressed in a given cell type. Transcriptional regulation in eukaryotes depends largely upon how genes are packaged in chromatin. Eukaryotic cells contain a number of multiprotein complexes capable of remodelling chromatin through chemical modification of histone residues and/or through ATP hydrolysis to shift nucleosomes relative to the DNA sequence (Hargreaves and Crabtree 2011; Narlikar et al. 2013; Chen and Dent 2014). The activities of these complexes play important roles in the control of gene expression, DNA repair, and genome integrity (Clapier and Cairns 2009). Early studies of chromatin remodelling complexes classified several as co-repressors, based largely upon their component enzymatic activities, and how they could be shown to impact expression of reporter genes in vitro or in vivo (Wolffe 1997; Knoepfler and Eisenman 1999; Ahringer 2000). More recently, genome-wide analyses have revealed that the function of so-called co-repressor complexes is not as simple as once presumed: they are often found at sites of active transcription, and are required for transcriptional activation of some genes and repression of others (Reynolds et al. 2013). This raises the question of how a multiprotein complex, with defined enzymatic activities, can impart different regulatory functions with varying genetic context.

To understand how these co-repressors operate at the molecular level, we must first understand how the biochemical activities of complexes work to influence gene expression patterns. It is clear that certain histone modifications correlate with transcriptional status (Strahl and Allis 2000). For example, acetylation of histone H3 at lysine 27 (H3K27Ac) correlates strongly with active promoters and enhancers genome-wide, while trimethylation of H3K27 is associated with transcriptional inactivity. Yet it has recently been shown that polycomb repressive complex 2 (PRC2), responsible for methylating H3K27, rarely if ever acts to silence gene expression in mouse ES cells but rather acts to maintain gene silencing (Riising et al. 2014). Similarly, it is not clear whether histone deacetylase activity normally instructs the repression of promoters and enhancers or instead acts, like PRC2, to maintain or reinforce their inactivation (Henikoff and Shilatifard 2011). While nucleosome remodelling activity may be considered to “open or close” chromatin structure in a general way, exactly how a change in nucleosome density might result in either inhibitory or stimulatory effects on transcriptional activity of RNA polymerase II is not often clear.

Transcription factors drive developmental decisions, often exerting influence via enhancer sequences (Spitz and Furlong 2012). Active enhancers are brought into close proximity to their target promoters, thereby allowing bound transcription factors to interact with the RNA Polymerase machinery, frequently through the Mediator Complex (Kornberg 2005; Vernimmen and Bickmore 2015). Mediator influences transcription in many ways, including stimulating phosphorylation of the RNA Polymerase II C-terminal heptad repeat (Kim et al. 1994; Kornberg 2005; Allen and Taatjes 2015), which affects both transcription initiation and elongation (Buratowski 2009). Thus, the modulation of chromatin structure at enhancers or promoters by chromatin remodelling proteins could impact RNA Polymerase activity through a number of direct or indirect mechanisms.

The Nucleosome Remodelling and Deacetylation (NuRD) complex is an abundant, highly conserved multiprotein chromatin remodeller initially defined as a transcriptional repressor (Wade et al. 1998; Xue et al. 1998; Zhang et al. 1998). NuRD activity facilitates cell fate transitions in a range of different organisms and developmental contexts (Ahringer 2000; Chen and Dent 2014; Laugesen and Helin 2014; Signolet and Hendrich 2015). NuRD combines class I lysine deacetylase activity, encoded by the Hdac1 and 2 proteins, with the Swi/Snf-type ATPase/nucleosome remodelling activity of Chd4. Also in the mammalian complex are the histone chaperone proteins Rbbp4/7, at least one of the SANT-domain proteins Mta1, 2, or 3; zinc finger proteins Gatad2a or −b; the Cdk2ap1 protein and the scaffold protein Mbd3 (Kloet et al. 2015). In mouse ES cells, NuRD activity modulates the transcription of pluripotency-associated genes, maintaining expression within a range that allows cells to effectively respond to differentiation signals (Reynolds et al. 2012a). The NuRD complex associates with virtually all active enhancers and promoters in mouse ES cells (Miller et al. 2016). This suggests that NuRD activity contributes a previously unrecognised component of active transcription.

Whether just one or both of the two distinct enzymatic activities of NuRD controls transcriptional modulation has not been determined. NuRD activity stimulates loss of H3K27 acetylation, providing a substrate for PRC2-mediated trimethylation at some genes (Reynolds et al. 2012b), a function that may fulfil either an instructive or reinforcing role. Conversely, the nucleosome remodelling activity of Chd4 has been shown to generally increase nucleosome density at target sites and thus facilitate lineage commitment through control of gene expression probability (Moshkin et al. 2012; Morris et al. 2014; O'Shaughnessy-Kirwan et al. 2015; de Dieuleveult et al. 2016), but exactly how nucleosome remodelling mechanistically impacts gene expression is unknown. A much more detailed understanding of the interplay between these two activities is required to understand how transcription is so precisely controlled in developmental contexts.

In this study we set out to determine how NuRD function impacts the transcription machinery to modulate gene expression levels. Using an inducible system with fine temporal resolution, we show that transcriptional control by the NuRD complex is initially exerted through its chromatin remodelling activity. NuRD acts predominantly at enhancers to maintain nucleosome density, resulting in reduced association of Mediator and fine control of RNA Polymerase II C-Terminal Domain (CTD) phosphorylation. This work defines the mechanism of action of an abundant chromatin remodelling complex with high temporal resolution and defines the molecular events underlying its control of transcriptional regulation in mammalian cells.

## Results

### NuRD associates with sites of active transcription

As a first step towards understanding NuRD activity in ES cells, we mapped chromatin binding sites genome-wide for two component proteins, Chd4 and Mbd3, in mouse ES cells using ChIP-seq. Mbd3 ChIP-seq using a polyclonal antibody against Mbd3 or against an epitope tag (Avi-3xFLAG) that was knocked-in to the endogenous *Mbd3* locus (Supplemental Fig. S1A) gave essentially identical results. The presence of this short C-terminal tag on the Mbd3 protein had no detectable adverse effects on the ability of Mbd3 to interact with NuRD components (Supplemental Fig. S1B) or to respond to differentiation conditions (Supplemental Fig. S1C). Mbd3-A3xFLAG cells were used to derive transgenic mice by blastocyst injection, and the resulting line was bred to homozygosity, thus confirming that the C-terminal tag did not detectably interfere with Mbd3 function.

Both Chd4 and Mbd3 were found to associate with chromatin extensively in ES cells (Fig. 1A), as has been reported in somatic cells (Miccio et al. 2010; Zhang et al. 2012; Gunther et al. 2013; Shimbo et al. 2013; Morris et al. 2014; Schwickert et al. 2014). Mbd3 binding was almost completely coincident with Chd4 binding, consistent with Mbd3 functioning exclusively within the NuRD complex (Fig. 1A). In contrast, more than twice as many Chd4-bound peaks as Mbd3-bound peaks were identified. This is consistent with both NuRD-dependent and NuRD-independent functions for the Chd4 protein (Williams et al. 2004; O'Shaughnessy and Hendrich 2013; O'Shaughnessy-Kirwan et al. 2015). By plotting ChIP-seq enrichment at co-bound (NuRD peaks) and Chd4-only peaks (Fig. 1B), we find that Mbd3 is nonetheless detected at many Chd4-bound loci, albeit at varying levels. We therefore conclude that at approximately 45% of all Chd4-bound sites, Chd4 and Mbd3 show a similar degree of enrichment (which we refer to as “NuRD-bound” sites) whereas the remaining Chd4-bound sites show reduced, more variable Mbd3 enrichment, possibly indicating that they are less frequently occupied simultaneously by Chd4 and Mbd3.

**Figure 1.**
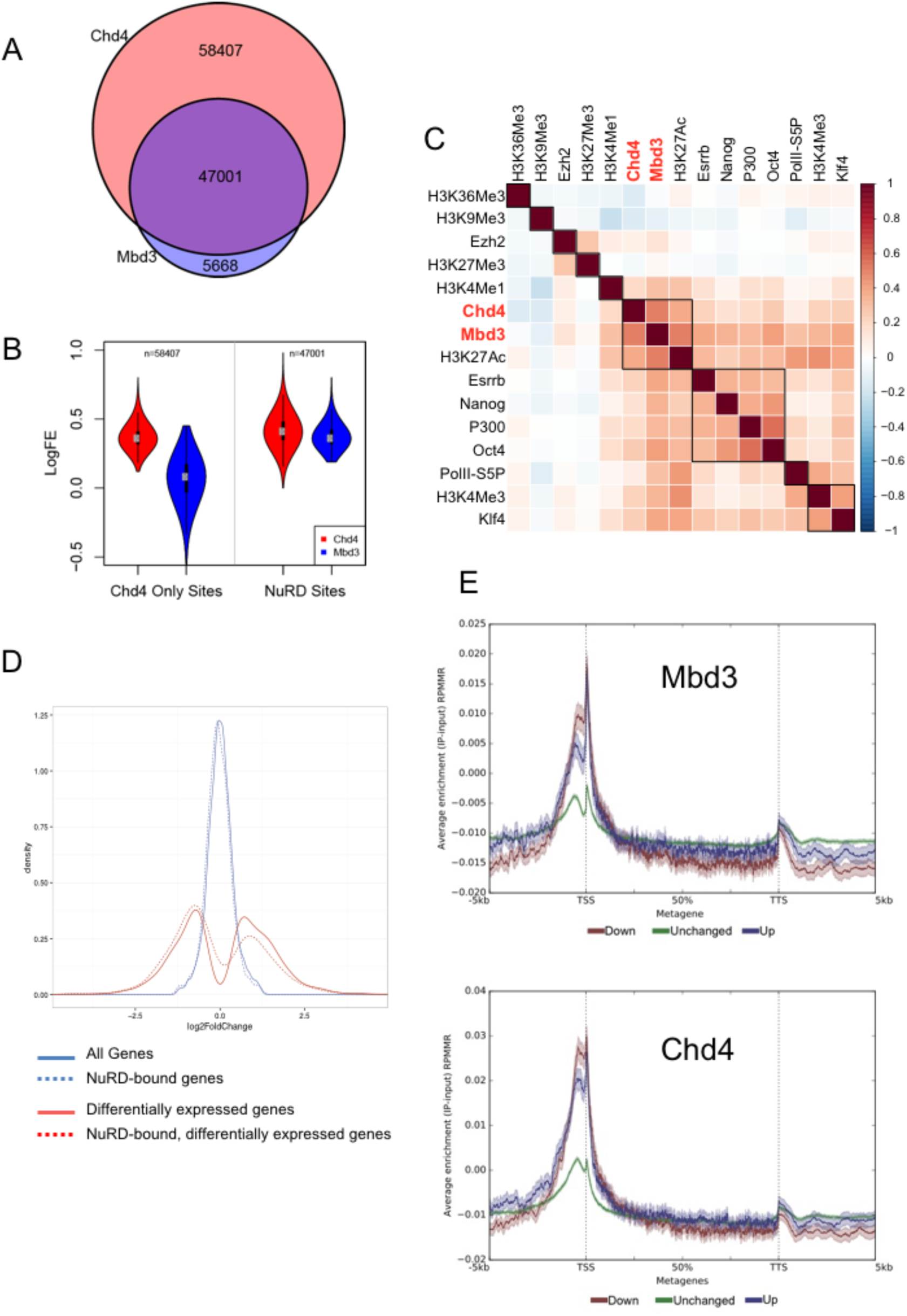
NuRD modulates active transcription. A. Mbd3- or Chd4-bound peaks (blue and pink circles, respectively) as identified by ChIP-seq in mouse ES cells. Numbers of peaks corresponding to Chd4-only, Mbd3 and Chd4, or Mbd3-only are indicated. B. Chd4 (red) or Mbd3 (blue) ChIP-seq (log_10_ fold enrichment) for sites exclusively bound by Chd4 (left) and those co-bound by Mbd3 and Chd4 (NuRD Sites; right). Numbers of peaks in each category are indicated. C. Correlation between ChIP peaks for the indicated histone modifications and transcription factors in 2i/LIF. Boxes indicate the highest correlations. Datasets used are listed in the supplementary Materials and Methods. D. Changes in gene expression in *Mbd3*-null vs wild-type ES cells. Red lines indicate genes that exhibit a significant change in mutant cells, blue lines indicate no significant change. Genes showing a ChIP-seq peak for Mbd3 and Chd4 between -2 Kb and + 0.5 Kb of the annotated TSS are indicated as dotted lines. y-axis: kernel density estimation; x-axis: log_2_ fold change (KO/WT). E. Mbd3 and Chd4 ChIP-seq density ±5 Kb across a metagene. Mean enrichment levels ± 95% confidence intervals are plotted for genes showing no change (green), a reduction (red) or increase (blue) in expression in *Mbd3*-null ES cells. See also Figure S1.

Hierarchical clustering of Chd4 and Mbd3 ChIP-seq datasets with those from published ChIP-seq data compiled in CODEX (Sanchez-Castillo et al. 2015) shows that binding for both proteins correlates strongly with indicators of active promoters and enhancers such as H3K27Ac, H3K4Me1, H3K4Me3, P300 and the initiating form of RNA Polymerase II (PolII-S5P; Fig. 1C). NuRD association strongly correlates with the patterns observed for the pluripotency-associated transcription factors Oct4, Nanog, Esrrb and Klf4. In contrast, NuRD component binding is anti-correlated with a mark of silent chromatin (H3K9Me3) and with one deposited across gene bodies (H3K36Me3). Weak correlation is seen for Ezh2 and trimethylated H3K27, consistent with NuRD cooperation with PRC2 at a subset of binding sites in ES cells (Reynolds et al. 2012b; Signolet and Hendrich 2015). These data show that NuRD is found predominantly at sites associated with transcription initiation, i.e. active enhancers and promoters, in ES cells.

NuRD function has been shown to influence gene expression both positively and negatively, indicating that NuRD performs a more complex role in transcription than mere silencing (Reynolds et al. 2012a). Consistent with this assertion, we identify a similar number of genes with increased or decreased expression in *Mbd3*^-/-^ ES cells (Fig. 1D). Enrichment of both Mbd3 and Chd4 is slightly increased at genes that display a decrease in expression level in *Mbd3*-null ES cells compared to those showing increased levels, but enrichment at both classes of genes is much greater than that observed at loci where no changes in transcription are detected (Fig. 1E). We therefore conclude that NuRD is a general regulator of active transcription in ES cells.

### NuRD activity rapidly induces gene expression changes

To better understand how NuRD function impacts gene expression we sought to observe the effect of acute NuRD recruitment to chromatin. To achieve this we took advantage of a system that allowed us to restore NuRD activity to a cell in which it is lacking, and then monitor the impact on transcription over time. Briefly, we used Mbd3-null ES cells in which we express the “b” isoform of the Mbd3 protein (Hendrich and Bird 1998) fused to mouse estrogen receptor domains at both the N- and C-termini. In the absence of tamoxifen, Mbd3 is confined to the cytoplasm and the ES cells adopt an Mbd3-null phenotype lacking functional NuRD complex (Reynolds et al. 2012b). Upon tamoxifen addition Mbd3 protein translocates into the nucleus and NuRD assembly is enabled (Fig. 2A). Mbd3 became detectable in the nucleus ≤15 minutes after tamoxifen addition, where it could be found to interact with endogenous Chd4, indicating NuRD complex formation (Figs 2A, B).

**Figure 2.**
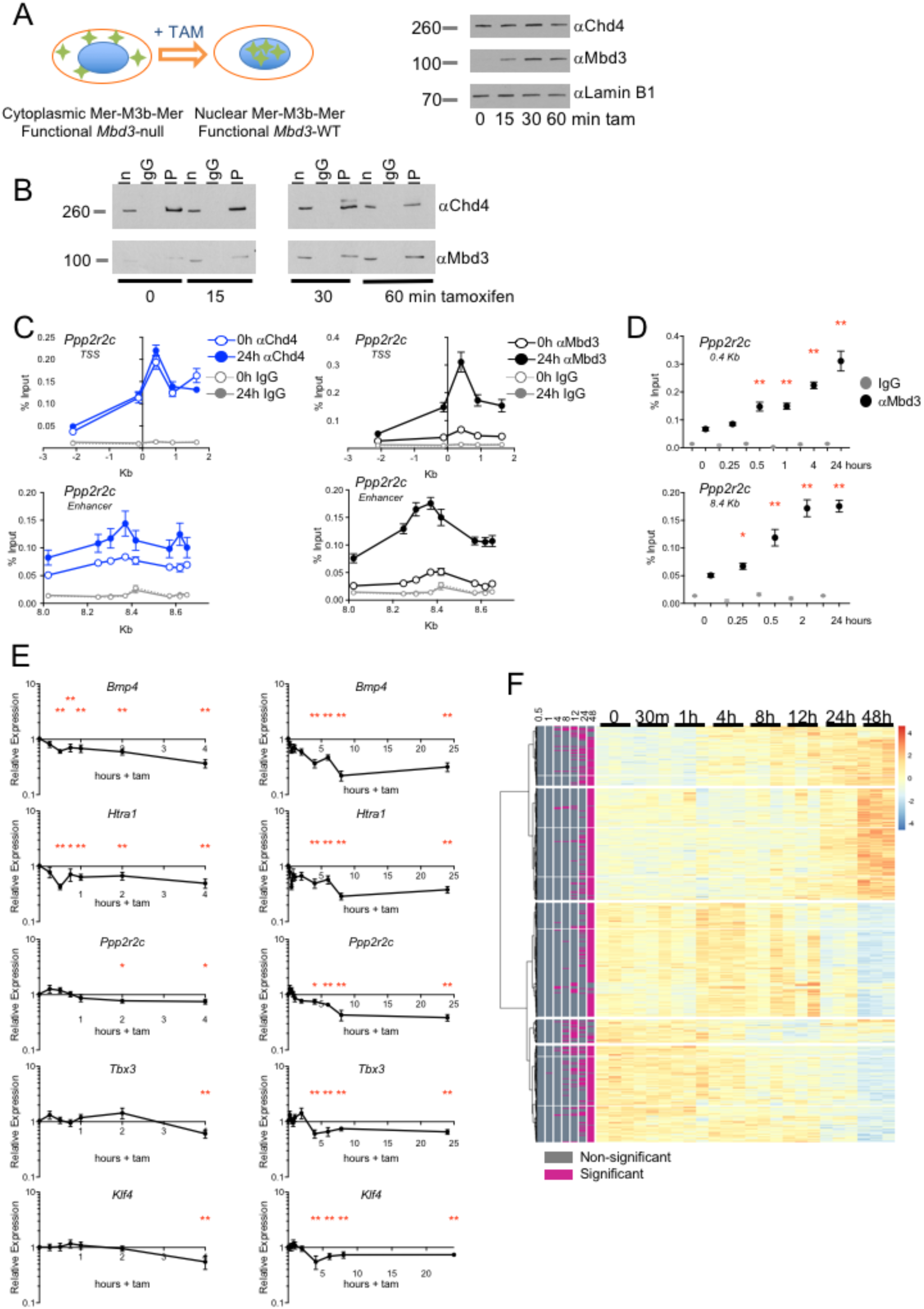
An Mbd3 induction system restores NuRD activity to *Mbd3*-null ES cells. A. Left: Model of the induction system: *Mbd3* null ES cells (left) contain Mer-Mbd3b-Mer (green diamonds, Mer-M3b-Mer) in the cytoplasm. Upon addition of tamoxifen Mer-Mbd3b-Mer enters the nucleus (blue circle). Right: Nuclear extracts were probed for Chd4, Mbd3 and Lamin B1 (as a loading control) at indicated times after tamoxifen addition. Mer-M3b-Mer is detected in the nucleus within 15 minutes of tamoxifen treatment by an anti-Mbd3 antibody and reaches its maximum level after 30 minutes. B. Nuclear Mer-M3b-Mer is rapidly incorporated into the NuRD complex. Chd4 was immunoprecipitated from nuclear protein extracts following tamoxifen exposure for indicated times and probed for the presence of Mbd3. In = 10% input, IgG = IgG control, IP = Chd4 immunoprecipitation. C. Induced Mbd3 is detected at both promoters and enhancers of target genes by chromatin immunoprecipitation (ChIP). ChIP-qPCR was carried out for Chd4 (blue line), Mbd3 (black line), and an IgG control (grey line) across the promoter and an enhancer of known NuRD target gene *Ppp2r2c* at 0 and 24 hours of tamoxifen treatment. N ≥ 9. See also Figure S2A. D. Mbd3 recruitment occurs rapidly, within 30 minutes of induction. Enrichment relative to input for the peak of the Mbd3 ChIP signal in panel C was plotted across a time course of tamoxifen addition. Significant enrichment of Mbd3 relative to no tamoxifen occurs from 30 minutes onwards (** P<0.001, * P<0.01). N ≥ 9. E. qRT-PCR data (mean of relative expression ± SEM; plotted relative to time 0) for nascent RNA of indicated genes over 24 hours of the time course of tamoxifen addition. Panels on the left show only the first four hours of data displayed at right. Asterisks indicate points at which expression is significantly different from that at time 0 using a two-tailed *t*-test (* P≤0.01, **P≤ 0.001). In panels to the right asterisks indicate only the 4, 6, 8, and 24-hour time points. N ≥ 9. F. Unsupervised clustering of gene expression changes over indicated times of the time course of tamoxifen exposure as measured by RNA-seq. Data for each replicate are shown. Genes exhibiting a significant change at 1, 4, 8 or 24 hours are shaded pink in the first, second, third, or fourth column, respectively.

Mbd3 was detectable on promoter and enhancer chromatin by ChIP-qPCR between 15 and 30 minutes after tamoxifen addition, and enrichment increased over the first 24 hours (Figs 2C, 2D; Supplemental Fig. S2A). Levels of Chd4 enrichment at some, but not all Mbd3 target loci were increased during this time course, and in steady-state conditions the presence of Mbd3 protein correlated with an increase in Chd4 enrichment at Mbd3-bound regions (Supplemental Fig. S2B). This indicates that while Chd4 is present at target loci in the absence of Mbd3, the presence of Mbd3 protein may stabilise Chd4 on some chromatin targets. The Mbd3-inducible system thus provides a means to monitor the direct effects of NuRD formation at its target sites in ES cells.

Restoration of NuRD activity had a rapid impact on expression of genes known to be repressed by the complex. Some genes, such as *Htra1* and *Bmp4*, showed reduced levels of nascent RNA from 30 minutes after Mbd3 induction (Fig. 2E). Other genes, such as *Ppp2r2c*, *Tbx3* and *Klf4*, responded more slowly, exhibiting a reduction in nascent RNA 2-4 hours after tamoxifen addition (Fig. 2E). Changes in transcription were specific to NuRD formation as a mock treatment of the cells with ethanol (the solvent for tamoxifen) did not significantly alter expression levels (Supplemental Fig S2C). Persistent and significant changes in steady-state mRNA levels were first detectable by total RNA-seq four to eight hours post-tamoxifen addition which increased steadily through 48 hours (Fig. 2F). The timescale for induction of NuRD complex formation, recruitment to chromatin and subsequent changes in expression therefore provides a means to probe the molecular changes that underlie NuRD dependent transcriptional regulation.

### Histone H3 lysine 27 acetylation is uncoupled from NuRD-dependent gene changes in expression

NuRD enzymatic activities include protein deacetylation. As we had previously shown that acetylation of histone H3 lysine 27 (H3K27Ac) is anticorrelated with the presence of NuRD activity (Reynolds et al. 2012b), we first asked whether induction of NuRD had any immediate impact on local or global levels of H3K27 acetylation. Surprisingly, given that both *Ppp2r2c* and *Htra1* showed reduced transcription within four hours of tamoxifen addition, levels of H3K27Ac as measured by ChIP-qPCR relative to total H3 levels were unchanged 24 hours after NuRD formation at the promoters of either gene, but decreased by 48 hours (Fig. 3A). At finer temporal resolution, H3K27Ac levels at both promoters showed slight initial increases within four hours post tamoxifen exposure (though to a statistically significant extent only at *Htra1*), followed by a gradual decrease through 48 hours (Fig. 3B). A similar pattern was seen for another marker of active promoters, trimethylation of histone H3 lysine 4 (H3K4Me3; Figs 3A, B).

ChIP-seq across the time course revealed no consistent change in levels of H3K27Ac at the promoters of genes where rapid reductions in nascent RNA levels were observed (Fig. 3C). The promoters of genes responding to Mbd3 induction with significant changes in mRNA levels similarly showed no perturbation of H3K27ac (Fig. 3D) or H3K4Me3 (Supplemental Fig. S3) marks. As acute changes in gene expression could be driven by activity at enhancers, in addition to promoters, we also assessed H3K27Ac levels at NuRD-bound enhancers over the time course, but again could detect no significant alterations to H3K27 acetylation (Fig. 3E). Although there was a slight reduction in signal for both H3K27Ac and H3K4Me3 at NuRD-bound sites after four hours, initial levels were restored by 24 hours (Fig. 3E, Supplemental Fig S3B). Together, these results indicate that levels of H3K27Ac at enhancers and promoters, and of H3K4Me3 at promoters follow, rather than instruct, changes in transcription that occur in response to introduction of NuRD activity in ES cells.

**Figure 3.**
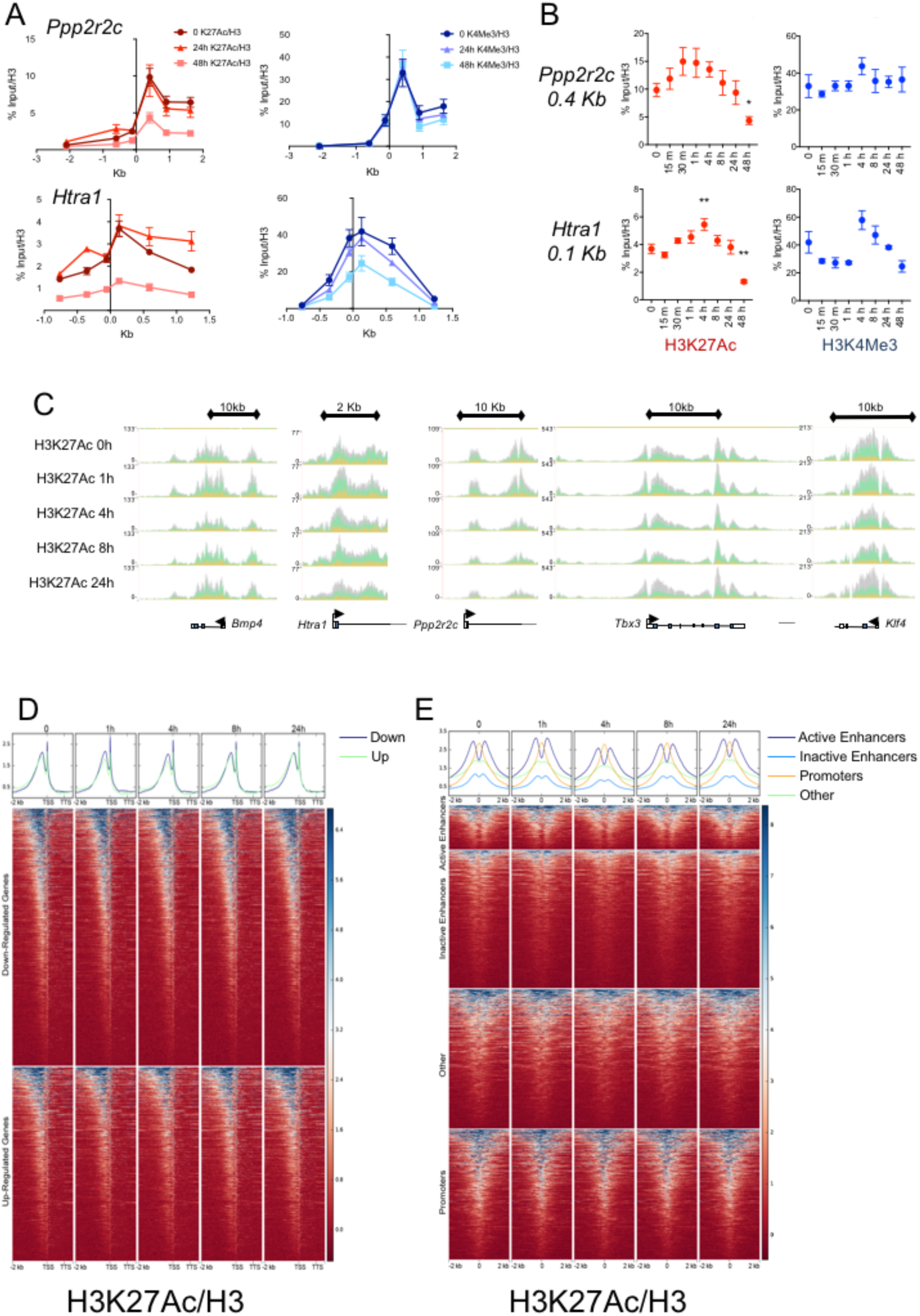
H3K27Ac and H3K4Me3 levels do not immediately correlate with gene expression changes. A. ChIP-qPCR for H3K27Ac (red) and H3K4Me3 (blue) are plotted relative to H3 ChIP across the *Ppp2r2c* and *Htra1* transcription start site for 0, 24, and 48 hours following tamoxifen addition. Mean ± SEM are plotted. N ≥ 9 B. ChIP for H3K27Ac (red) and H3K4Me3 (blue) at the peak of enrichment from panel A is displayed across the time course of tamoxifen exposure. Mean ± SEM. (** P<0.01, * P<0.05 relative to 0 hours using a two-tailed *t*-test). N ≥ 6. C. ChIP-seq traces visualised on the UCSC Genome Browser for the *Ppp2r2c*, *Tbx3* and *Bmp4* loci across the Mbd3 induction time course. The schematics below show the gene organisation and direction of transcription. Replicate traces are stacked in the display (H3K27Ac: N=3; H3K4Me3: N=2). D. ChIP-seq for H3K27Ac/H3 for each time point of Mbd3 induction across genes showing decreased (top panels) or increased (lower panels) expression by RNAseq during the first 48 hours of tamoxifen exposure. Genes are plotted as a metagene including 2Kb upstream of the transcription start site (TSS). TTS refers to the polyA-addition site. Mean signal for genes showing decreased (blue) or increased (green) expression is plotted above. E. ChIP-seq for H3K27Ac/H3 for each time point of Mbd3 induction centred at peaks of NuRD binding ± 2 Kb, classified into active and inactive enhancers, promoters, or other sequences. Mean signal for each category is shown above.

### NuRD remodels positioned nucleosomes in enhancers

In addition to histone deacetylase activity, NuRD contains nucleosome remodelling activity encoded by Chd4. Assessing chromatin structure globally by MNase-seq in *Mbd3*-null ES cells and wild-type controls, we identified a striking difference in nucleosome positioning adjacent to Mbd3 binding sites (Fig. 4A, Supplemental Fig S4A). While Mbd3 association tended to occur in nucleosome-depleted regions, sequences immediately adjacent to the binding sites displayed increased MNase protection in Mbd3-null ES cells, consistent with the presence of a highly positioned nucleosome (Fig. 4A, Supplemental Fig S4A). This effect was most pronounced at active enhancers, but was also apparent at inactive enhancers and promoters bound by Mbd3 (Fig. 4A). Notably, the effect was specific to Mbd3-bound sites, and was much less pronounced at all Chd4-bound sites or loci containing a Chd4 peak but not an Mbd3 peak. This is surprising given that the remodelling activity is conferred by the Chd4 protein, and perhaps suggests that co-localisation of Mbd3 affects not only the stability of Chd4 association with chromatin but also its activity.

In order to probe NuRD’s chromatin remodelling specificity more generally, nucleosome density in wild-type and *Mbd3*-null ES cells was mapped relative to a range of transcription factor bound sites derived from previously published ChIP-seq data (Fig. 4B, Supplemental Fig S4B). This analysis revealed pronounced changes in occupancy of one to two nucleosomes adjacent to binding sites for enhancer-associated transcription factors (Klf4, Nanog, Esrrb, Oct4) as well as general transcriptional activators (Med12, P300), but less so for the binding sites of a promoter-associated protein (Tbp) (Fig. 4B). No change in nucleosome positioning was evident adjacent to Ctcf binding sites (Fig. 4B). Loci bound by Ctcf showed a very slight increase in MNase protection in *Mbd3*-null ES cells, though considerably less than that reported for cells lacking the NuRF or SNF2H chromatin remodellers (Bohla et al. 2014; Qiu et al. 2015; Kwon et al. 2016; Wiechens et al. 2016). This indicates that the nucleosome remodelling activity of NuRD acts at enhancers to normalise nucleosome density across regions of relatively open chromatin.

**Figure 4.**
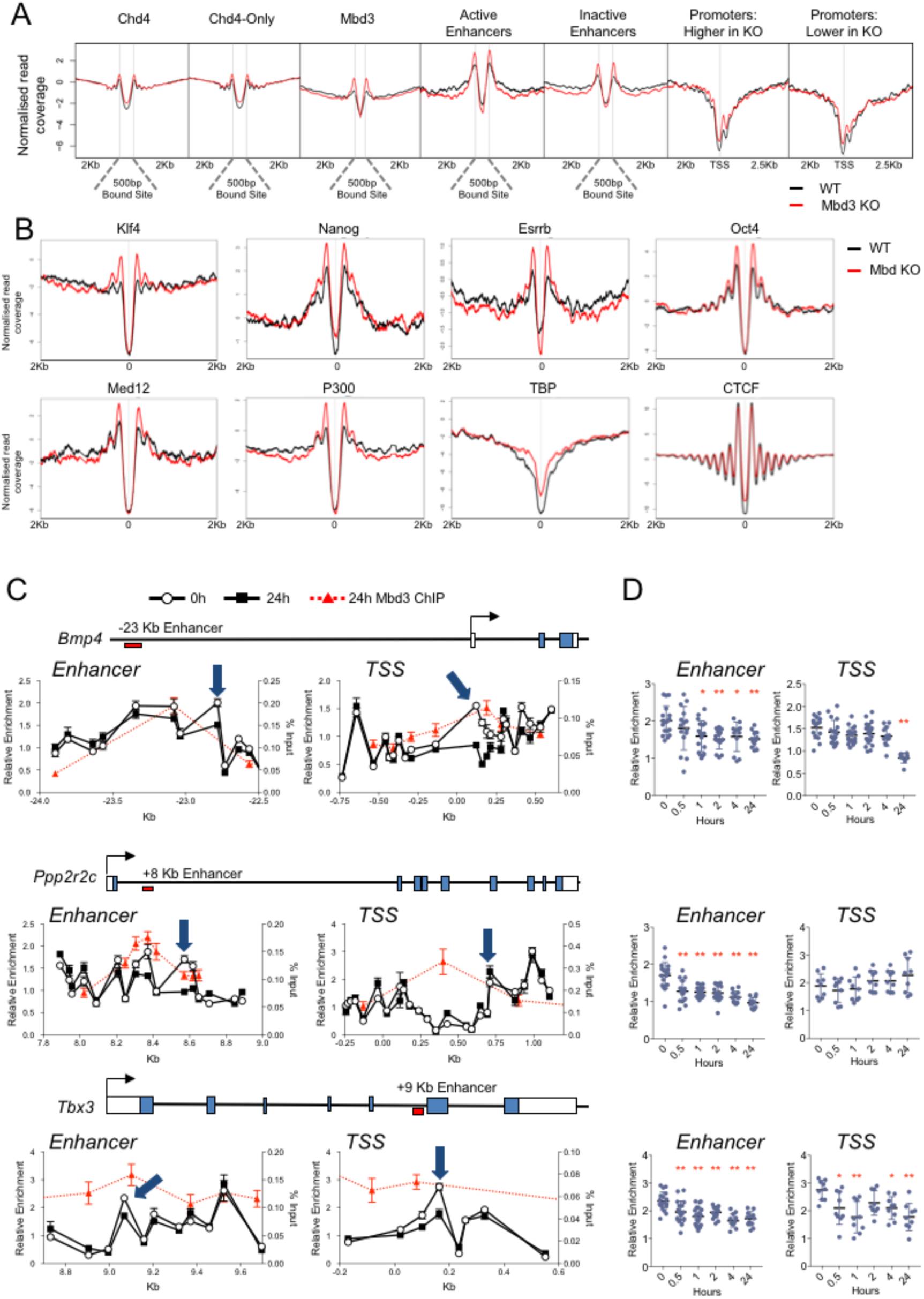
NuRD maintains nucleosome density across enhancers. A. MNase-seq data from wild-type (black) and *Mbd3*-null (red) ES cells plotted across Mbd3-binding sites (scaled to 500bp), including 2 kb of flanking sequence for all sites, active and inactive enhancers. Also plotted is MNase-seq for 2 Kb on either side of annotated transcription start sites (TSS) of genes showing higher or lower expression in *Mbd3* KO ES cells as compared to wild-type controls. Similar analyses from a replicate experiment are shown in Figure S4. B. MNase-seq data as in panel C centred at the ChIP-seq peaks defined for the indicated published datasets. See also Figure S4 and Table S4. C. MNase-qPCR data (mean ±SEM) for the 0 and 24h time points plotted across an enhancer or the TSS for indicated genes. The x-axis indicates Kb relative to the annotated TSS for each gene. Overlaid in red is ChIP-qPCR data for Mbd3-ER at 24 hours post tamoxifen addition. The blue arrow indicates the position further analysed in panel D. A schematic of each gene is shown above the ChIP-qPCR panels. Filled and open boxes represent coding and non-coding exons, respectively, and the red box below the line indicates the position of the relevant enhancer. See also Figures S4C and S5. N ≥ 9. D. MNase-qPCR data across the time course of tamoxifen exposure for the position indicated by a blue arrow in panel C. Blue circles represent individual data points, with mean ± SD indicated with black lines (** P<0.01, * P<0.05 relative to 0 hours). N ≥ 9.

To examine more precisely how NuRD remodels enhancer chromatin, we next monitored nucleosome positions across the Mbd3 induction time course by MNase-qPCR at genes repressed by NuRD within 30 minutes (*Bmp4*), 2 hours (*Pppr2c*) or 4 hours (*Tbx3*) of tamoxifen exposure. Positioned nucleosomes could be detected in *Mbd3*-null ES cells at both enhancers and promoters of all three NuRD-responsive genes (Fig. 4C, Supplemental Fig S4C). Consistent with our steady-state MNase-seq data, Mbd3 induction resulted in rapid (30-60 minutes) loss of positioning of specific nucleosomes within enhancers associated with all three genes (Figs 4C, D). This very rapid change coincided with, or preceded a change in transcription, and persisted for the duration of the time course. At all three enhancers examined the nucleosome being displaced was that immediately adjacent to the peak of Mbd3 binding (Fig. 4C), again consistent with our steady-state analysis. Sequences bound by nucleosomes further from the peak of NuRD binding showed little or no change in MNase accessibility, indicating that this is not a general effect across the enhancer but rather is specific to individual nucleosomes. NuRD-dependent changes in nucleosome positioning were also identified at *Bmp4* and *Tbx3* promoters, but only in the case of *Tbx3* was the effect detectable prior to transcriptional response (Fig. 4C). Together these data indicate that NuRD functions to displace positioned nucleosomes specifically within target enhancer sequences.

### NuRD activity modulates the protein binding repertoire of enhancers

What effect might such a precise change in nucleosome positioning have at active enhancers? Mechanistically, enhancers act as binding sites for transcription factors to exert influence on the activity of RNA Polymerase II (recently reviewed in (Engel et al. 2016)). Mbd3 and Chd4 genome-wide binding patterns are highly correlated with those of pluripotency-associated transcription factors (Fig. 1B). NuRD-bound sites show a modest increase in the steady-state ChIP-seq signal for Nanog in Mbd3-null cells, but not for Klf4 (Fig. 5A). To assess the effect of NuRD function on transcription factor binding globally we performed ChIP-seq for Klf4 and Nanog across the Mbd3 induction time course. While there appears to be an initial loss of Klf4 enrichment at all classes of Mbd3-bound sites 30 minutes after tamoxifen addition, levels then recover by four and 24 hours, before statistically significant expression changes are widespread. It seems likely that such a transient loss is not directly linked to transcriptional regulation. Nanog enrichment showed a gradual increase in average binding over the time course (Fig. 5B). No change in the levels of nuclear Nanog or Klf4 protein was detected across the time course (Supplemental Fig S6A). This demonstrates that NuRD activity can impact the ability of transcription factors to bind to their targets, but the nature of this influence appears to differ for specific factors. Notably, mean Nanog signal was not decreased at 24 hours relative to time 0 (i.e. as the cells become more like wild-type), indicating that the steady-state increase in Nanog signal observed in Mbd3-null ES cells may be a longer-term consequence of loss of NuRD function.

**Figure 5.**
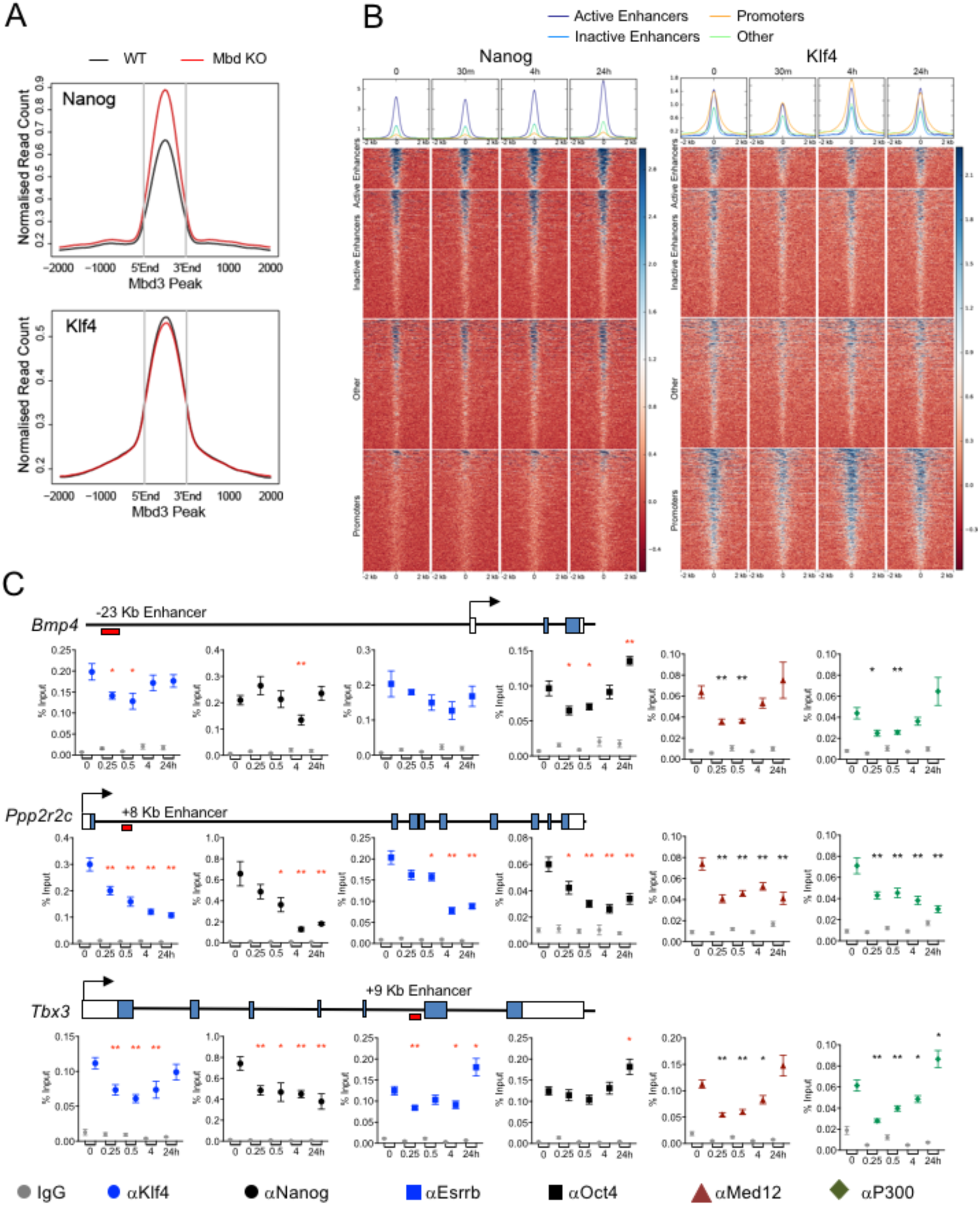
NuRD does not prevent transcription factor access to chromatin. A. ChIP-seq profiles for Nanog and Klf4 across scaled Mbd3 peaks in wild-type (black) or *Mbd3*-null (red) ES cells. B. ChIP-seq for Nanog (left) and Klf4 (right) at indicated times following Mbd3 induction across NuRD peaks that coincide with active or inactive enhancers, promoters, or at other sequences, centred at the peak of NuRD binding. Plots at the top show mean enrichment for active enhancers (dark blue), inactive enhancers (light blue), promoters (orange) or other sequences (green) for each time point. See also Figure S6. C. ChIP-qPCR for Klf4 (blue circles), Nanog (black circles), Esrrb (blue squares), Oct4 (black squares), Med12 (maroon triangles) P300 (green diamonds) and IgG control (grey) at the peak of binding (see Figure S6B) at indicated enhancers across the time course of tamoxifen exposure. A schematic of each gene is shown above the ChIP-qPCR panels, as in Figure 4. Mean ± SEM is plotted for each point; time in hours is indicated along the x-axis. ** P<0.01, * P<0.05 relative to 0 hours. N > 3 for all points. See also Figure S5. N ≥ 9.

To examine the relationship between NuRD-dependent nucleosome remodelling and transcription factor binding in greater detail, we next explored how NuRD-dependent nucleosome remodelling impacts transcription factor occupancy at the *Bmp4*, *Ppp2r2c* and *Tbx3* enhancers by ChIP-qPCR. Here we extended the transcription factors assayed to include Esrrb and Oct4, which similarly show no change in nuclear protein levels over the time course (Supplemental Fig S6A). Reconstitution of the NuRD complex resulted in rapid loss of Klf4 binding to all three enhancers tested with kinetics similar to that of nucleosome displacement, although binding is regained at the *Bmp4* and *Tbx3* enhancers by 24 hours (Fig. 5C). Nanog binding was also rapidly lost upon Mbd3 induction, and remained absent at both the *Ppp2r2c* and *Tbx3* enhancers, but not at the *Bmp4* enhancer. Esrrb and Oct4 displayed variable responses to Mbd3 induction: both were lost from the *Ppp2r2c* enhancer, but at the other enhancers no consistent pattern was observed, with any initial loss replaced by a gain of enrichment by 24 hours (Fig. 5C, Supplemental Figs S6A, B). Together these data do not support a global, general role for NuRD activity in either preventing or enhancing the binding of sequence-specific transcription factors to their enhancer targets. Rather, while induction of NuRD activity does influence transcription factor association with enhancer chromatin, this results in transient and/or locus-specific changes that varied between the four sequence-specific transcription factors profiled.

In addition to serving as binding platforms for sequence-specific transcription factors, enhancers are also associated with general transcriptional activators such as Mediator or P300 (Calo and Wysocka 2013). In contrast to the sequence-specific factors, the binding of both a component of the Mediator complex (Med12) and P300 was reduced at all three enhancers immediately after Mbd3 induction (Fig. 5C). This was not associated with any changes in levels of these two proteins during the induction time course (Supplemental Fig S6A). At the *Ppp2r2c* and *Tbx3* enhancers Med12 remained depleted over the 24-hour duration, while at the *Bmp4* enhancer binding of both Med12 and P300 was regained after an initial pronounced reduction (Fig. 5C). The very rapid time scale of Med12 and P300 loss as NuRD localises to enhancers indicates that either NuRD itself, or NuRD-mediated nucleosome remodelling directly interferes with the ability of Med12 and P300 to associate with enhancers.

### Chromatin-mediated inhibition of Mediator binding induces a rapid loss of RNA Polymerase II CTD phosphorylation

Mediator can promote the activity of TFIIK to phosphorylate the C-terminal heptad repeat of RNA Polymerase II at the serine 5 position (Kim et al. 1994; Robinson et al. 2016). Levels of RNA polymerase phosphorylated at the S5 position (S5P) were rapidly depleted from the TSS regions of *Bmp4, Ppp2r2c* and *Tbx3* upon NuRD formation (Fig. 6A), coincident with a reduction in Mediator association. Loss of S5P signal was particularly rapid at the *Ppp2r2c* and *Tbx3* genes (≤30 minutes), which preceded any change in nascent transcript levels (Fig. 2E) and by 24 hours correlated with a reduction in total RNA polymerase ChIP signal (Fig. 6A). Similarly, ChIP-seq revealed a general reduction in S5P levels at the TSS of all genes showing altered expression within the first four hours of NuRD induction (Fig. 6B).

**Figure 6.**
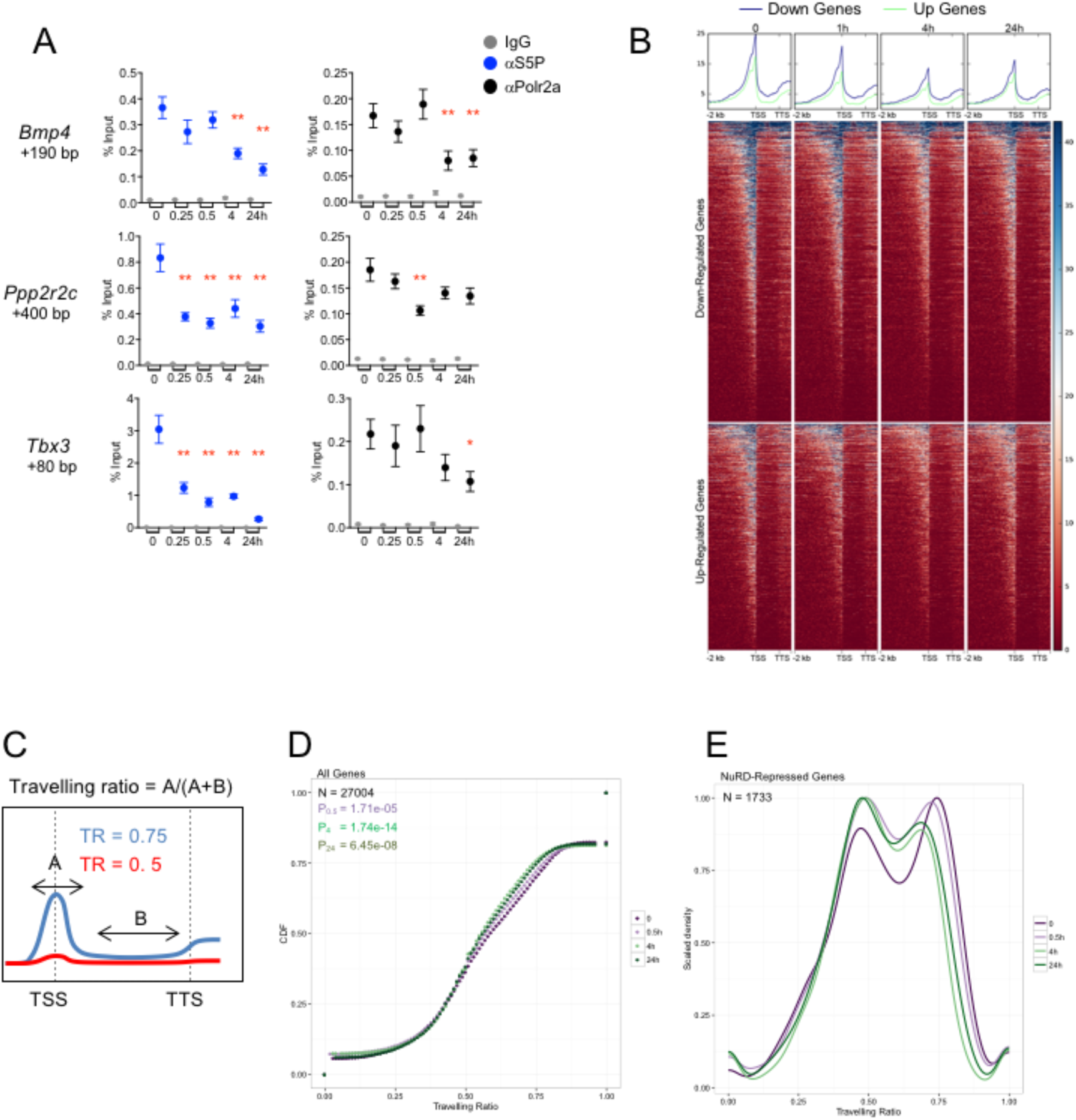
RNA Polymerase II rapidly and globally responds to NuRD complex formation. A. ChIP-qPCR for S5P (blue), total RNA Polymerase II (Polr2a; black) and IgG control (grey) at the indicated position relative to the TSS for each gene is plotted across the tamoxifen addition time course; Mean ± SEM is plotted (** P<0.01, * P<0.05 relative to 0 hours). N > 3 for all points. N ≥ 9. B. ChIP-seq signal for S5P is plotted across down (blue)- and up (green)-regulated metagenes as in Figure 3D. Plots at the top show mean enrichment at each time point. C. Schematic of travelling ratio calculation. The blue line represents S5P ChIP-seq signal across a paused gene (TR = 0.75), whereas the red line shows signal across an unpaused gene (TR = 0.5). TSS = transcription start site, TTS = transcription termination site. D. Cumulative travelling ratio calculated from S5P ChIP-seq data at time 0 (dark purple), 30 minutes (0.5H; light purple), 4 hours (light green) and 24 hours (dark green). CDF = Cumulative Density Function. E. Travelling ratio plotted against scaled density for genes showing decreased or expression during the first 24 hours of tamoxifen exposure.

RNA polymerase II levels are often displayed as a travelling ratio (Adelman and Lis 2012) (Fig. 6C). This gives an indication of the relative abundance of S5P at promoters versus the gene body. It can be used to indicate the degree of transcriptional pausing, although initiation rate and elongation rate also influence travelling ratio (Ehrensberger et al. 2013). ChIP-seq for the S5P form of RNA polymerase II showed that introduction of NuRD activity resulted in a rapid, global decrease in travelling ratio, detectable as early as 30 minutes after induction, and which persisted beyond 24 hours (Fig. 6D).

NuRD-repressed genes could be subdivided into two main groups: those with a travelling ratio of about 0.5, and those around 0.75 (Figs 6C, E). A travelling ratio of 0.5 represents an essentially flat profile, with no enrichment at the TSS, while 0.75 indicates accumulation of S5P around the TSS (Fig. 6C). Induction of NuRD activity results in both a reduction in the number of genes within the high travelling ratio group (height of the peak at TR = 0.75, Fig. 6E) as well as a decrease in the average travelling ratio in this group (leftward shift of the peak at TR = 0.75; Fig. 6E). The majority of genes repressed by NuRD showed a high travelling ratio prior to induction, but rapidly shifted to the lower travelling ratio upon NuRD induction (Fig. 6E). As NuRD can be found at virtually all active enhancers and promoters in ES cells, this is consistent with a rapid and global influence of NuRD activity on RNA Polymerase II phosphorylation.

### NuRD limits Mediator recruitment to enhancers during developmental transitions

NuRD activity facilitates exit from the self-renewing state in ES cells through control of gene expression (Reynolds et al. 2012a). We therefore investigated whether NuRD restricts Mediator access to enhancer sequences as ES cells exit the self-renewing state. We chose to examine changes in enhancer chromatin of ES cells after 24 hours in differentiation conditions, as this is the point immediately preceding a change in the kinetics of silencing of the naïve pluripotency genes *Klf4* and *Zfp42* (*Rex1*), and activation of the primed pluripotency marker *Otx2* between wild-type and *Mbd3*-null ES cells (Fig. 7A). Enhancers associated with *Klf4*, *Zfp42* and *Otx2* were bound by Mbd3 in both self-renewing and differentiation conditions, and all loci featured increased nucleosome positioning in the absence of NuRD activity 24 hours after switching to differentiation conditions (Figs 7B, 7C). During this window we also detected an increase in Mbd3 association at these enhancers in wild-type cells, further supporting a role for NuRD in modulating nucleosome occupancy in tandem with transcriptional regulation (Fig. 7B). Furthermore, this increased nucleosome positioning is accompanied by increased association of two different Mediator components with these enhancers in *Mbd3*-null cells (Med1 and Med12, Fig. 7D), verifying that NuRD activity restricts both nucleosome positioning and Mediator access to enhancers during developmental transitions.

**Figure 7.**
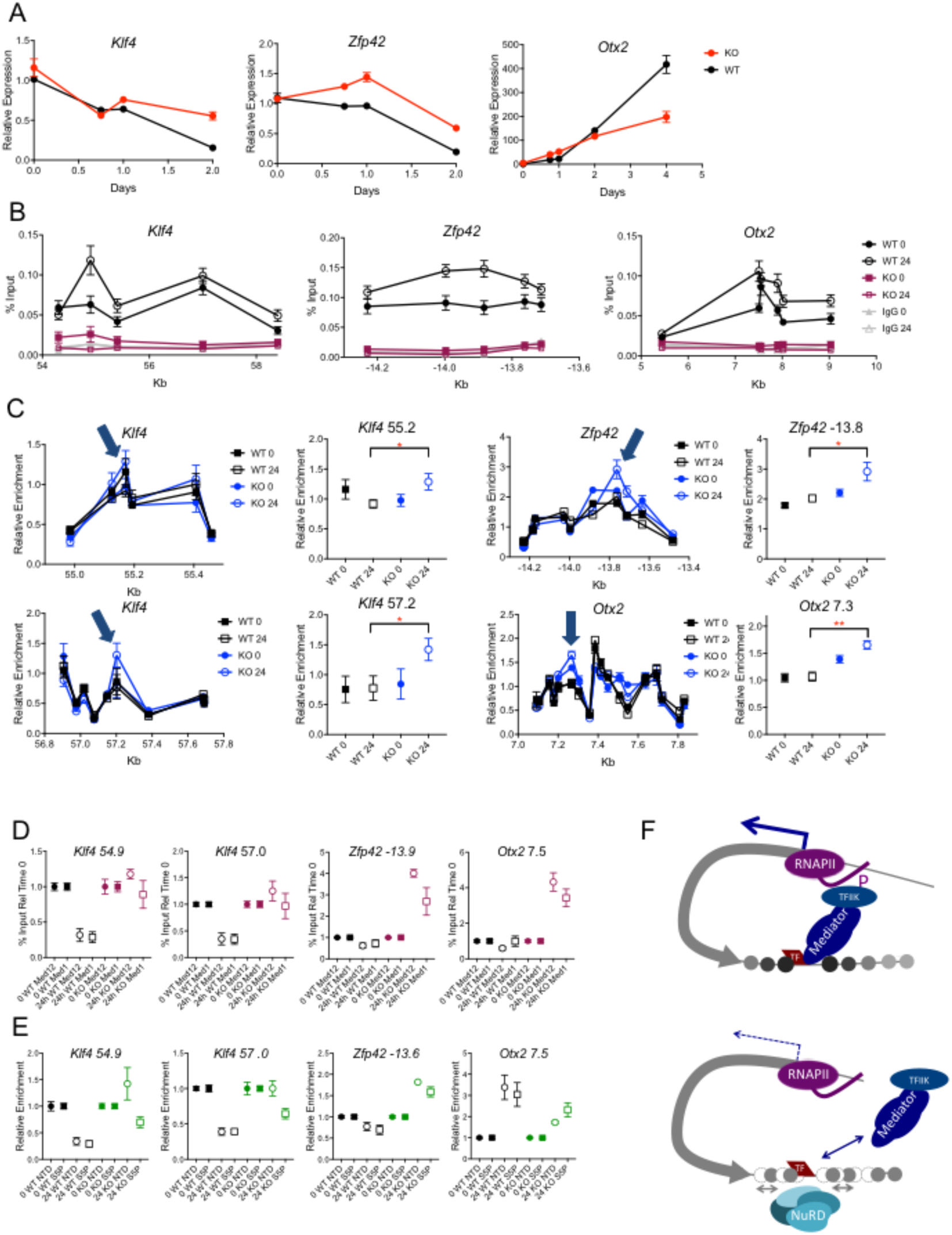
NuRD-mediated remodelling of enhancer chromatin facilitates Mediator access during lineage commitment. A. Expression of *Klf4*, *Zfp42* and *Otx2* over a differentiation time course relative to that in wild-type cells in 2iL conditions. Mean ± SEM are plotted for all points. N ≥ 9. B. ChIP-qPCR for IgG (grey) and Mbd3-FLAG in wild-type (black) and Mbd3-null (magenta) ES cells across indicated enhancer sequences in 2iL (0) or after 24 hours in differentiation conditions (24). Mean ± SEM are plotted for all points. N ≥ 12. C. MNase-qPCR profiles across two enhancers associated with the *Klf4* gene and one each for the *Zfp42* and *Otx2* genes are plotted for wild-type (black) or Mbd3-null ES cells (blue) in 2iL conditions (0) and after 24 hours in differentiation conditions (24). Data for the points indicated with blue arrows are plotted at right. Asterisks indicate a significant difference between WT and KO at the 24h time point (** P<0.01, * P<0.05). Mean ± SEM are plotted for all points. N ≥ 9. D. ChIP-qPCR for Med12 (circles) or Med1 (squares) in wild-type (black) or Mbd3-null ES cells (magenta) before and after 24 hours in differentiation media. Data are plotted relative to levels at time 0. Mean ± SEM are plotted for all points. N ≥ 6. E. ChIP-qPCR for total RNA Polymerase II (circles) or S5P (squares) in wild-type (black) or Mbd3-null ES cells (green) before and after 24 hours in differentiation media. Data are plotted relative to levels at time 0. Mean ± SEM are plotted for all points. N ≥ 9. F. Model of NuRD-mediated transcriptional regulation. In the absence of functional NuRD complex (top) nucleosomes at enhancers are highly positioned. Mediator is able to bind readily to this region and stimulates TFIIH to phosphorylate serine 5 of the RNA Polymerase II CTD at the transcription start site. Serine 5 phosphorylation leads to an increase in transcriptional activity. In wild type-cells (bottom) NuRD activity increases nucleosome dynamics, particularly at enhancers. This reduces the interaction of Mediator complex with the enhancer region thereby reducing the activity of the TFIIK subunit of TFIIH on RNA Polymerase. By restricting the interaction between Mediator and the enhancer in this way NuRD acts to modulate levels of gene expression rather than acting as a switch to simply turn transcription on or off.

In wild-type cells the enrichment of RNA Polymerase associated with these enhancers after 24 hours of differentiation correlates with transcriptional change: we detected reduced association with the *Klf4* and *Zfp42* enhancers, but an increase at the *Otx2* enhancer (Fig. 7E). While at this much lower time resolution (days vs hours) and a more heterogeneous system than the Mbd3 induction time course we did not detect the rapid change in RNA polymerase II serine 5 phosphorylation (Fig. 6), we found that RNA polymerase status was dysregulated in the absence of Mbd3 under these conditions. In *Mbd3* null cells the *Zfp42* enhancer showed elevated levels of RNA Pol II association at 24 hours, consistent with the observed increase in *Zfp42* transcription relative to undifferentiated cells (Fig. 7E). The *Klf4* enhancers also displayed increased RNA Pol II at 24 hours, which preceded a silencing failure detected at 48 hours (Figs 7A, E). Similarly, reduced association with RNA polymerase II was observed at the *Otx2* enhancer in *Mbd3* null cells at 24 hours despite showing increased Mediator binding, and this was immediately followed by an activation defect in *Otx2* transcription (Figs 7A, E). Thus NuRD-dependent changes in nucleosome positioning, Mediator binding and RNA Polymerase II association at enhancer sequences all precede major changes in gene regulation, consistent with them playing a causative role in transcriptional change. We conclude that transcriptional control of a subset of genes associated with induction of differentiation requires appropriate modulation of chromatin structure by the NuRD complex, which in turn restricts Mediator access to enhancers to accurately modulate transcription (Fig. 7F).

## Discussion

Although many chromatin remodelling proteins are essential for mammalian development, the actual mechanics of how they impact transcription in vivo remains ill-defined. Here we show that NuRD, an abundant chromatin remodelling complex found at virtually all sites of active transcription in ES cells, exerts a transcriptional modulatory function by restricting the abundance of serine 5 phosphorylated RNA Pol II. The primary consequence of NuRD recruitment to enhancers is homogenisation of nucleosome density which limits access of transcriptional activators such as Mediator and P300 to enhancer sequences, possibly by altering the transcription factor binding repertoire. This, in turn, limits the abundance of the S5P form of RNA polymerase II, which ultimately impacts transcription levels (Fig. 7F). We therefore propose that enhancer chromatin structure, in the form of nucleosome positioning, is a primary determinant of transcriptional output.

The influence of enhancers on the transcription machinery has long been defined in general terms: enhancers bring transcription factors to the vicinity of RNA polymerase II, providing a “positive” or “activating” influence. Decades of functional and structural work has recently culminated in a refined model of how Mediator, which is believed to bridge enhancers and the transcription machinery, directly interacts with the CTD of RNA Polymerase II to facilitate its phosphorylation by the TFIIK subunit of TFIIH (Plaschka et al. 2016; Robinson et al. 2016). By limiting Mediator access to enhancers, NuRD buffers the stimulatory influence of Mediator upon TFIIK activity (Fig. 7F). At the majority of transcribed sequences this buffering has no detectable influence on steady-state expression levels (Fig. 1D). At a subset of genes this buffering results in reduced transcription, while at others transcription is increased. Though initially counterintuitive, the opposing effects of this increased buffering and subsequent decrease in S5P levels may be similar to that of RNA Polymerase pausing, which has been shown to impact transcriptional output in a similar way in both *Drosophila* development and in mouse ES cells (Gilchrist et al. 2008; Henriques et al. 2013).

Mediator is believed to be recruited to enhancers through association with transcription factors (Spitz and Furlong 2012; Allen and Taatjes 2015). NuRD activity was found to exert a variable influence on the binding profiles of Klf4, Nanog, Esrrb and Oct4, and it is likely that other transcription factors are also sensitive to NuRD activity. This scenario would be consistent with a recent study in which knockdown of *Mbd3* was found to result in decreased nucleosome abundance at various transcription factor binding sites, which was interpreted as NuRD controlling transcription factor occupancy (Hainer and Fazzio 2015). Subtle changes in the protein binding syntax of an enhancer can have a large influence on the activity of the target gene (Sokolik et al. 2015; Farley et al. 2016; White et al. 2016), and it is likely that Mediator and P300 are sensitive to this. Adding transcription factor binding and/or enhancer interactions to the recent Mediator/RNA Polymerase II structural models will likely help to better understand how Mediator interacts with enhancer chromatin (Plaschka et al. 2016; Robinson et al. 2016).

A separation of nucleosome remodelling and deacetylase activities was predicted in the initial reports describing the NuRD complex (Tong et al. 1998; Wade et al. 1998; Xue et al. 1998; Zhang et al. 1998). The in vivo targets of the lysine deacetylase activity of NuRD are unclear, although anticorrelation of H3K27Ac levels with NuRD function has been observed in steady-state conditions (Reynolds et al. 2012b). In vitro, the histone deacetylase components of the NuRD complex show little substrate specificity (Feller et al. 2015; Zhang et al. 2016). The extent to which NuRD deacetylates non-histone proteins, and what effects such activities might impart on the transcription process, remain to be determined. In our system perturbation of H3K27Ac marks is not associated with the early stages of transcriptional regulation, but rather follows expression level changes (Fig. 3). This observation is consistent with studies in yeast and flies reporting that transcription induction in some contexts does not require covalent histone modifications, and contributes to the discussion about the role of histone modifications in transcription (Henikoff and Shilatifard 2011; Zhang et al. 2014; Perez-Lluch et al. 2015). It is possible that NuRD-directed H3K27 deacetylase activity functions predominantly to reinforce gene expression programs. The synergistic interaction between NuRD and PRC2 is most consistent with this maintenance role for NuRD-associated deacetylase activity (Morey et al. 2008; Reynolds et al. 2012b; Riising et al. 2014). In agreement with this notion, there is evidence that NuRD-dependent histone deacetylation contributes to longer-term enhancer inactivation as ES cells are induced to differentiate (Whyte et al. 2012).

The molecular details of chromatin remodelling complex function have been somewhat enigmatic in mammalian cells. Genetic experiments prove that most of these complexes are essential for cell viability and/or early stages of mammalian development, while biochemical data demonstrate that such complexes can move, remove, or re-position nucleosomes, and that this often impacts transcription (Hargreaves and Crabtree 2011; Narlikar et al. 2013). Yet details of exactly how these biochemical activities are used by the cell to enable developmental decisions have remained elusive. Activating complexes, such as Ino80 or the BAF complex, are thought to render chromatin generally more accessible and thereby facilitate access by the transcriptional machinery and/or transcriptional activators (Hargreaves and Crabtree 2011; Narlikar et al. 2013; Wang et al. 2014), although presumably this would also allow access to repressors. Co-repressor complexes, such as NuRD, are believed to carry out the opposite reaction. Our work provides molecular detail to this general scheme. By restricting access of Mediator to enhancer sequences NuRD indirectly controls the levels of the initiating form of RNA Polymerase II genome-wide to facilitate control of transcription.

## Materials and Methods

### Tissue culture

ES cells were cultured in 2i/LIF (2iL) media on gelatin-coated plates unless otherwise specified. Translocation of Mbd3b protein to the nucleus was induced with 4-hydroxytamoxifen added directly to the culture media to a final concentration of 0.4nM for varying times as indicated. Alkaline phosphatase assays were performed by plating 1000 cells into a 6-well dish and expanding for 4 days prior to staining according to the manufacturer’s protocol (Sigma). N = 3 to 12 wells per condition. Colonies were scored blind to genotype.

### ChIP, ChIP-seq and RNAseq

Chromatin immunoprecipitation (ChIP) was performed as described (Reynolds et al. 2012a). For details, and antibodies used, see Supplementary Methods. ChIP-seq libraries were prepared using the NEXTflex Rapid DNA-seq kit (Illumina) and sequenced at the CRUK Cambridge Institute Genomics Core facility (Cambridge, UK) on the Illumina platform. RNA-seq libraries were prepared using the NEXTflex Rapid Directional RNA-seq kit (Illumina) and sequenced as above. ChIP-seq and RNA-seq, and transcriptome analyses re described in the Supplementary Methods. Primers for nascent transcript detection were made to a combination of intronic and exonic sequences and are listed in Supplementary Table 2. All other gene-specific expression assays were carried out using gene specific TaqMan probes (Life Technologies). Sequencing datasets are listed in Table S3.

### MNase seq and MNase qPCR

For nucleosome positioning analyses, cells were grown in 2iL conditions (supplemented with tamoxifen as noted) for MNase qPCR and in serum/LIF conditions for MNase-seq. Cells were harvested, washed, lysed, and digested using standard methods. Locus-specific primers and positions relative to annotated transcription start sites are listed in Table S2. Details of MNase-seq processing are in the Supplemental Methods.

## Acknowledgments

We thank Rob Klose, Austin Smith, Ernest Laue, Kristian Helin, and BDH Group members for discussions and/or comments on the manuscript. We also thank Maike Paramor and Joaquin Martinez Herrera for NGS library preparation, and Sally Lees for cell culture support. MX was funded through an EMBL PhD studentship, EJ was funded through a BBSRC PhD studentship and JS was funded through an MRC PhD studentship. Funding in the BH lab was provided by a Wellcome Trust Senior Fellowship and the EU FP7 Integrated Project “4DCellFate.” Funding in the PB lab was provided by EMBL and BBSRC. BH and PB labs further benefit from core funding to the Cambridge Stem Cell Institute from the Wellcome Trust and Medical Research Council.

